# Genetic control of tracheid properties in Norway spruce wood

**DOI:** 10.1101/720904

**Authors:** J Baison, Linghua Zhou, Nils Forsberg, Tommy Mörling, Thomas Grahn, Lars Olsson, Bo Karlsson, Harry X Wu, Ewa J. Mellerowicz, Sven-Olof Lundqvist, María Rosario García-Gil

## Abstract

➢ Through the use of genome-wide association (GWAS) mapping it is possible to establish the genetic basis of phenotypic trait variation. Our GWAS study presents the first such an effort in Norway spruce (*Picea abies* (L). Karst.) for the traits related to wood tracheid characteristics.
➢ The study employed an exome capture genotyping approach that generated 178 101 high quality Single Nucleotide Polymorphisms (SNPs) from 40 018 probes within a population of 517 Norway spruce mother trees. We applied a LASSO based association mapping method using a functional multi-locus mapping approach that utilizes latent traits, with a stability selection probability method as the hypothesis testing approach to determine significant Quantitative Trait Loci (QTLs). Expression of the identified candidate genes was examined using publicly available spruce databases.
➢ The analysis have provided 31 loci and 26 mostly novel candidate genes, majority of which showing specific expression in wood-forming tissues or high ubiquitous expression, potentially controlling tracheids dimensions, their cell wall thickness and microfibril angle. Among most promising candidates, the analysis identified *Picea abies BIG GRAIN 2* (*PabBG2*) with predicted function in auxin transport and sensitivity, and *MA_373300g0010* - similar to wall-associated receptor kinases (WAKs), both associated to cell wall thickness.
➢ The results demonstrate feasibility of GWAS to identify novel candidate genes controlling industrially-relevant tracheid traits in Norway spruce. The presence of many traits with several significant QTLs supports the notion that the majority of these traits are polygenic in nature.

## Introduction

Norway spruce is considered to be one of the most multipurpose species. Its wood provides various solid wood products as well as pulp and paper products (Mäkinen *et al*., 2002). It is considered one of the best raw-materials for production of mechanical pulp for many types of paper grades (Varhimo & Tuovinen, 1999). The properties of the tracheids have large influences on the quality of the final products, and also on process economy and sustainability, for solid wood as well as fibre-based products. Tracheid morphology and cell wall structure influence the flexibility of wood and fibres, interactions among fibres, as well as the mechanical, physical and optical properties of the end-products (Brändström, 2002). Consequently, identifying the genetic background of different tracheid traits as a basis for breeding may bring benefits for both industry and society.

Various long-term breeding programmes for the species are already being pursued with the goal to identify genotypes with high productivity and wood quality (Hannrup *et al*., 2004). Wood density and microfibril angle (MFA) are key indicators of wood quality as they influence strength and dimensional stability of solid wood (Yang & Evans, 2003). However, combining productivity with wood quality in Norway spruce is problematic due to negative genetic correlations between these traits (Chen *et al*., 2014). One of the tools helping to understand these genetically complex variations in forest trees is the integration of extensive genetic and phenotypic data in order to discern the genetics underlying these traits (Neale & Ingvarsson, 2008; A. J. Eckert *et al*., 2009; Parchman *et al*., 2012). Hence, knowing the genetic control of these variations may lead to better control of the end products.

With genomic resources now available, a large array of molecular markers has been availed for the studying and understanding of complex traits. The majoritiy of these traits are known to be predominantly polygenic in nature, and affected by environmental effects, hence the need to utilize techniques that target the whole genome (Hall *et al*., 2010). The availability of such an array of genomic resources has led to the reliable identification of Quantitative Trait Loci QTLs, which in conifers are traditionally detected using suitable segregating populations such as, full- or half-sib progenies. More recently, GWAS, also known as Linkage Disequilibrium (LD) mapping, have been applied as an alternative approach to QTLs detection from traditional pedigree-based mapping studies. GWAS accounts for historical recombination events in the natural population as compared to those observed in a pedigree based QTL mapping (Neale & Savolainen, 2004). When confounding factors are taken into consideration, LD mapping provides greater resolution than pedigree studies, since it utilizes markers in strong LD with putative causative genomic region (Cardon & Bell, 2001).

Many coniferous species are characterized by an outcrossing mating system and large population sizes which lead to a rapid LD decay within the genomes and low inbreeding coefficient (Neale & Savolainen, 2004). However, rapid and heterogenous decay in conifer LD (Pavy *et al*., 2012) can be a source of concern as proximal markers can be completely unlinked and therefore offer no predictive power to the quantitative trait that may be residing physically close (Thavamanikumar *et al*., 2014). Together with LD heterogeneity, population structure (Larsson *et al*., 2013), epistasis and Genotype x Environment interactions (GxE) (Gupta *et al*., 2005) are factors that if not carefully controlled can negatively impact on QTL identification. The utilization of LD mapping in the dissection of genetic backgrounds underpinning complex traits has been shown in several systems, for example, complex solid wood properties in Norway spruce (Baison *et al*., 2019), white spruce (Beaulieu *et al*., 2011) and *Eucalyptus* (Thavamanikumar *et al*., 2014), and detecting genes underlying ecological adaptations in *Populus* (McKown *et al*., 2014). The dissection of these complex traits can benefit from the application of mathematical functions that account for the year-to-year variation across the annual growth rings. The development of mathematical methods for the analysis of dynamic data has made it possible to develop functional mapping approaches (Ma *et al*., 2002; Xing *et al*., 2012) that firstly model the phenotypes using curve-fitting methods and then utilize the parameters describing the curve (latent traits) for independent association analysis (Li *et al*., 2014; Camargo *et al*., 2018).

GWAS can also increase our knowledge on molecular processes controlling tracheid traits. Such traits are to a large extent determined by the genes acting during wood development (Mellerowicz *et al*., 2001; Plomion *et al*., 2001). Tracheid traits can also be regulated non-cell-autonomously by processes that take place in other organs and tissues. For example, the activity of shoot apical meristem determines the availability of auxin in the cambium and developing wood (Masuda, 1990; Farquharson, 2008), whereas the photosynthetic activity in the needles influences the availability of sucrose for wood biosynthesis (Wegrzyn *et al*., 2010). Therefore, combining the knowledge of candidate genes with their expression analysis will give more insights to the biological processes shaping tracheids.

The major goal of this study was to identify causative allelic effects of genomic regions contributing to wood tracheid traits using LD mapping on exome sequence capture data. Due to the large size of the Norway spruce genome (20 Gb) and its highly repetitive nature, it presents a challenge to use whole genome re-sequencing approaches for the development of molecular markers. Approaches aimed at reducing these genome complexities, especially by either eliminating or drastically reducing the repetitive sequences have been developed (Buell *et al*., 2014). These approaches are referred to as reduced representation approaches as there are used as proxies for whole genomic sequencing. In this study, we have used exome capture, aiming at maximizing the capture of exonic regions of the genome only, thereby increasing the coverage and depth of genic sequence in our variant detection study. The analysis provided 26 mostly novel candidate genes for regulation of various tracheid traits, which, along with their expression patterns, give new insights to the tracheid traits determination, and offer key markers for early genetic selection in Norway spruce breeding.

## Materials and Methods

### Association mapping population

The association mapping population, phenotypic data and statistical analysis are described in Chen *et al*., (2014) and Hayatgheibi *et al*., (2018). Briefly, the mapping population for the association mapping population constisted of two progeny trails established 1990 in Southern Sweden: (S21F9021146 aka F1146 (trial1) and S21F9021147 aka F1147 (trial2)), composed of 1373 and 1375 half-sib families. A randomized incomplete block design with single-tree plots was employed for both trials. From the trials, 524 families in 112 sampling stands were selected for use in the investigation of wood tracheid properties. Wood increment cores with diameter of 12 mm were collected at breast height (1.3 m) from six trees from each of the selected families of each trial. A total of 5618 trees were sampled: 2973 trees from trial F1146 and 2645 from F1147.

### Phenotypic data generation

The radial variations of growth, wood and tracheid attributes from pith to bark were analysed using the SilviScan instrument (Evans 1994; Evans 2006) at Innventia, now RISE Bioeconomy, Stockholm, Sweden. SilviScan is an instrument for efficient measurement of radial variations in a multitude of properties from the same sample with high spatial resolution. High precision sample strips from pith to bark were produced from the increment cores and automatically scanned for radial variations in cross-sectional tracheid widths with a video microscope, in wood density with X-ray transmission and in structural orientations with X-ray diffraction, combined with image analysis. From these data, information on radial variations of further traits were derived, such as wall thickness, coarseness and MFA of tracheids, and stiffness of wood (MOE). The locations of the annual rings were identifies, as well as of their compartments of earlywood (EW), transitionwood (TW) and latewood (LW), using the “20-80 density” definition (Lundqvist *et al*., 2018), established for use in different types of studies, such as (Kostiainen *et al*., 2009; Franceschini *et al*., 2012; Fries *et al*., 2014). Averages for all rings and their compartments were calculated for the traits and organised to be ready for use in continued genetic evaluations, such as the work on solid wood traits (Chen *et al*., 2014), on tracheid traits (Chen *et al*., 2016) and by (Baison *et al*., 2019). The traits addressed in the current work are listed in Table 1.

For MFA, central peak regression mathematical functions were fitted to describe the MFA variation from juvenile towards mature wood, using procedures presented by (Hayatgheibi *et al*., 2018), including also pre-processing of the data for removal of outliers. A threshold value of MFA 20° for the fitted curves was chosen to define the age when a limit of an inner core of wood with inferior timber properties occurred, the transition age MFA_TA_. The averages of MFA for wood inside and outside this limit were calculated, MFA_CORE_ and MFA_OUTER_. This provided 3 latent traits for MFA.

**Table 1.**
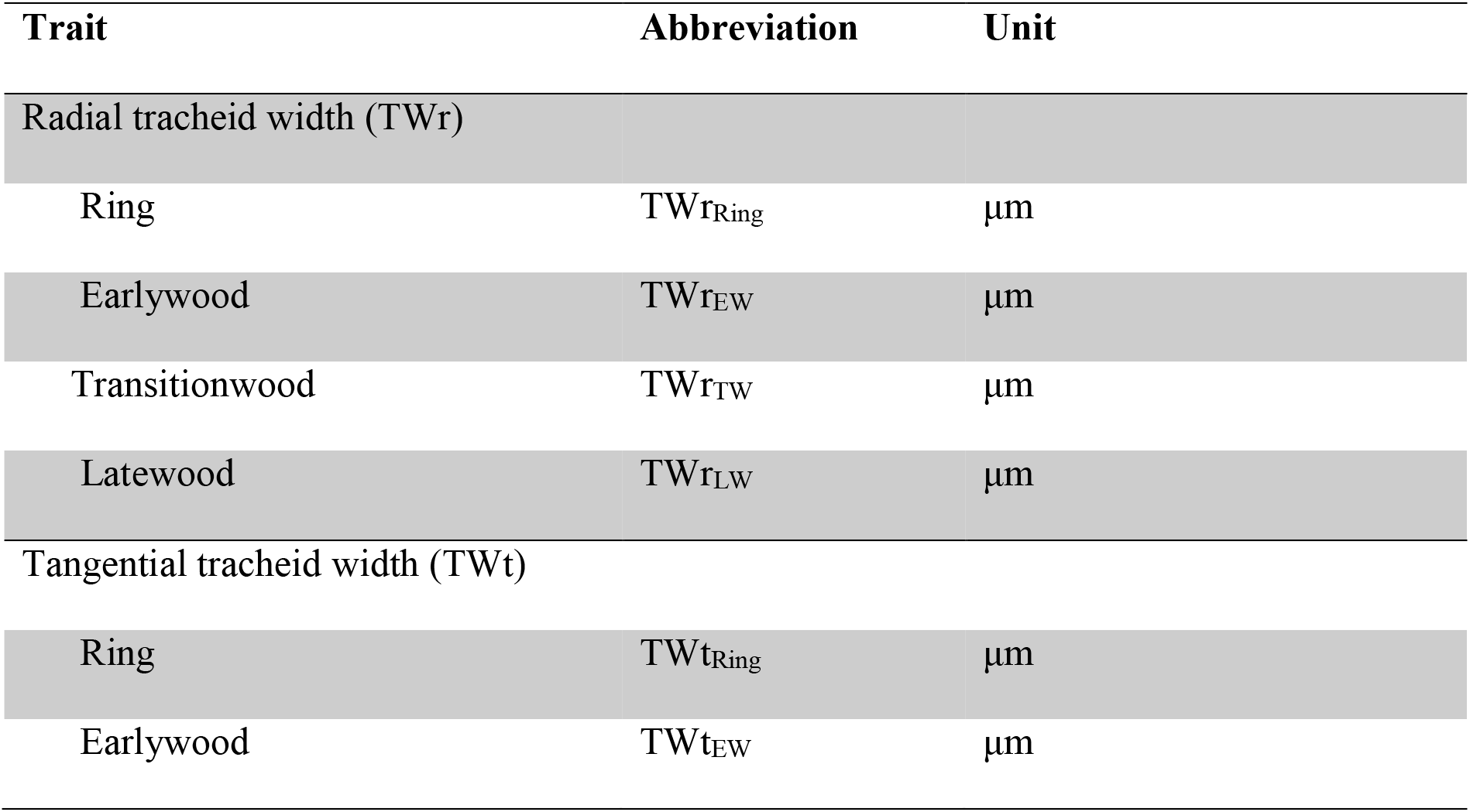
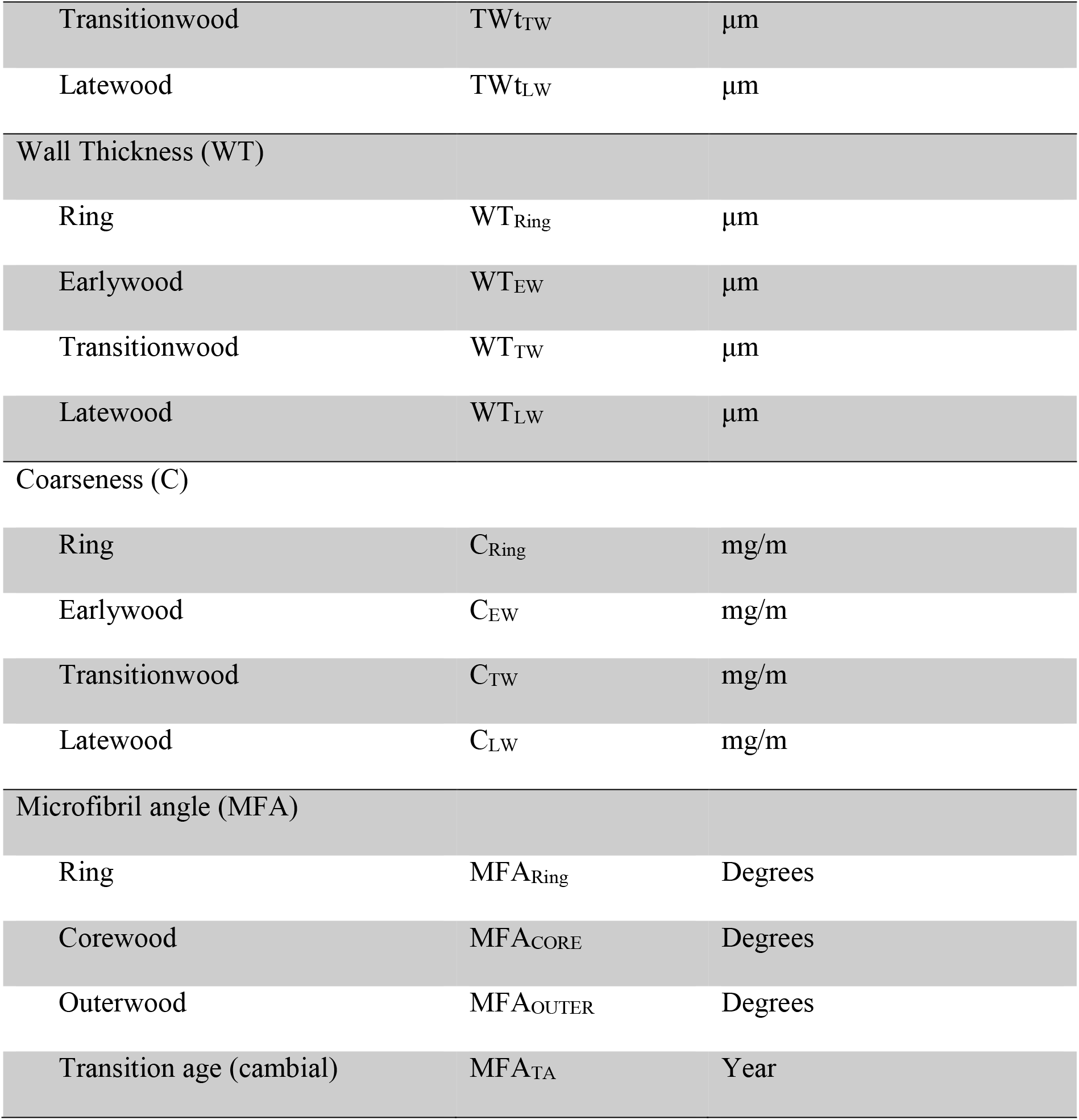
List of the traits, their abbreviations and measurement unit.

| Trait | Abbreviation | Unit |
| --- | --- | --- |
| Radial tracheid width (TW<sub>r</sub>) |  |  |
| Ring | TW<sub>rRing</sub> | µm |
| Earlywood | TW<sub>rEW</sub> | µm |
| Transitionwood | TW<sub>rTW</sub> | µm |
| Latewood | TW<sub>rLW</sub> | µm |
| Tangential tracheid width (TW<sub>t</sub>) |  |  |
| Ring | TW<sub>tRing</sub> | µm |
| Earlywood | TW<sub>tEW</sub> | µm |
| Transitionwood | TW<sub>tTW</sub> | μm |
| Latewood | TW<sub>tLW</sub> | μm |
| Wall Thickness (WT) |  |  |
| Ring | WT<sub>Ring</sub> | μm |
| Earlywood | WT<sub>EW</sub> | μm |
| Transitionwood | WT<sub>TW</sub> | μm |
| Latewood | WT<sub>LW</sub> | μm |
| Coarseness (C) |  |  |
| Ring | C<sub>Ring</sub> | mg/m |
| Earlywood | C<sub>EW</sub> | mg/m |
| Transitionwood | C<sub>TW</sub> | mg/m |
| Latewood | C<sub>LW</sub> | mg/m |
| Microfibril angle (MFA) |  |  |
| Ring | MFA<sub>Ring</sub> | Degrees |
| Corewood | MFA<sub>CORE</sub> | Degrees |
| Outerwood | MFA<sub>OUTER</sub> | Degrees |
| Transition age (cambial) | MFA<sub>TA</sub> | Year |

#### GWAS-LASSO

DNA extraction, variant detection and annotation and population structure on the genomic data utilized in this study was previously described by Baison et al (2019). Latent traits describing how the traits developed with age were calculated in two steps, see Baison et al (2019); posteriorly, association between latent traits and SNP genotypes (GWAS) was conducted with a LASSO penalized regression method as described by Li et al (2014).

In brief, estimated breeding values (EBV) were computed for each annual ring by cambial age to reduce site and block effects (see Chen et al 2014). In a second step, linear splines were applied to reconstruct time trajectories based on by annual ring EBV. Multiple knots, or latent traits, were then fitted to the EBV describing the shape of the time trajectories (Fig 1).

#### Candidate gene mining

To assess putative functionality of SNPs with significant associations, gene ontology and network analysis of putative genes and their associated orthologs was performed against the NorWood v1.0 database (http://norwood.congenie.org; (Jokipii-Lukkari *et al*., 2017)) hosted by ConGenIE (http://congenie.org/). After the identification of the QTL, the Norway spruce contigs linked to the significant SNPs were extracted from the web based database congenie (congenie.org/blast). The complete Norway spruce contigs that harboured the QTLs that were not annotated in the ConGenIE were used to perform a nucleotide BLAST (Blastn) search, using the option for only highly similar sequences (megablast) in the National Center for Biotechnology Information (NCBI) nucleotide collection database (https://blast.ncbi.nlm.nih.gov/Blast.cgi?).

### Results and Discussion

#### Trait trajectories

In order to visualize the nature of the trait variation over the successive years of wood development, the raw data for the traits were plotted for 19 consecutive cambial ages (Fig 1). For traits with complex growth trajectories the application of functional mapping enables the utilization of all the longitudinal data per trait, thus giving a well-informed analysis of temporal trends (Li *et al*., 2014).

The ring MFA initially decreased from an average across the trees of about 30° at the pith and stabilized after reaching a cambial age of about 10 years at an average of 10-12° (Fig 1a). The adapted central peak curves combined with the threshold at 20° resulted in an average of five years for MFA_TA_, defining the inner core of lower quality timber with AM performed for the latent traits of MFA_CORE_ and MFA_OUTER_.

For all the other tracheid phenotypes: wall thickness, radial tracheid width, tangential tracheid width and coarseness (Fig1), family means of β_0_ (intercept) and β_1_ to β_3_ (effects of knot 1 to 3, see Baison et al., 2019) were implemented in the association mapping. Candidate gene loci were identified only for the intercepts β_0_.

**Fig 1.**
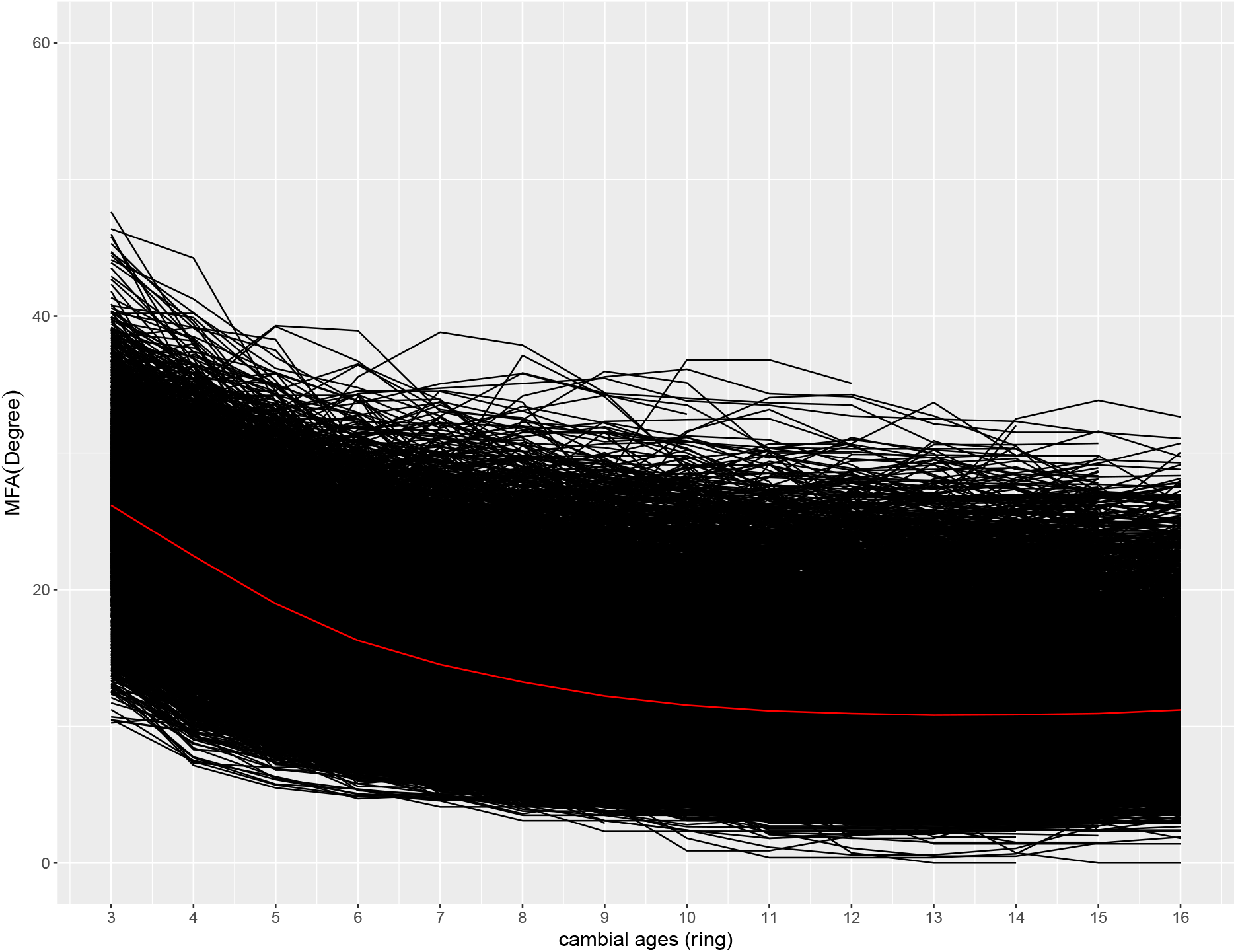
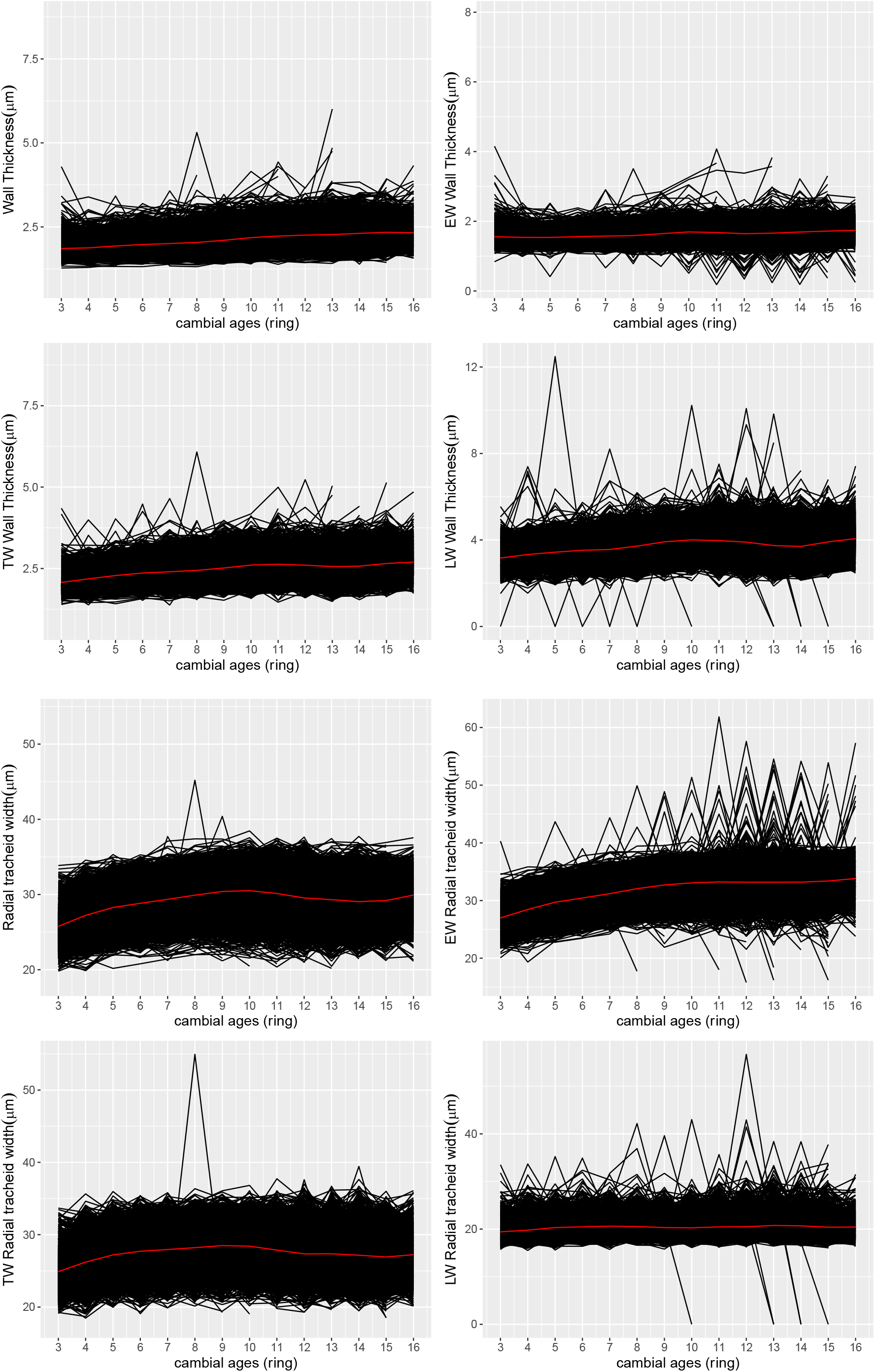
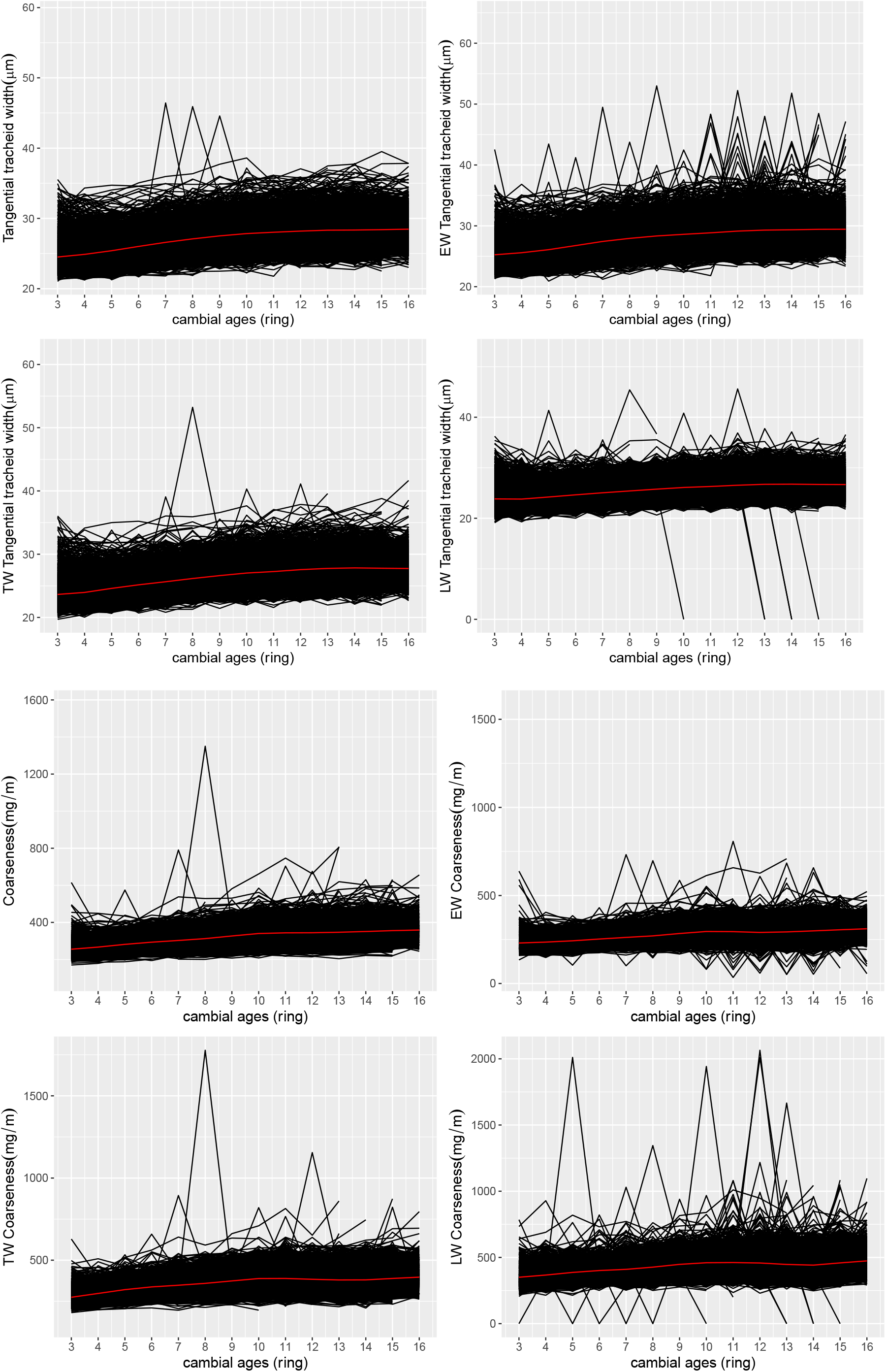
Trajectories of MFA, wall thickness, radial tracheid width, tangencial tracheid width, and coarseness related traits plotted against cambial age for individual trees. These individual trajectories were then used to determine which latent trait to be applied in the association mapping.

#### Genetic associations detected and modes of gene action

A total of 30 significant associations were detected across the 18 traits with fraction phenotypic variances being explained (PVE) ranging between 0.01 to 3.79% (Table 2), using an Stability selection probability (SSP) minimum inclusion frequency of 0.52. Seven of the 30 marker trait associations for which dominance and additive effects could be calculated were consistent with partially to fully dominant effects (0.50 < |*d/a*| < 1.25). The remaining 23 markers were all determined to have an additive (|*d/a*| < 0.50) modes of inheritance (Table 2). The relationships between the genotypic classes of markers associated to a phenotype were consistent with these patterns (Fig. 2). All the significant SNPs were located within upstream regions with only two SNPs (MA_10434903g0010_165836 and MA_11172g0010_12016) being within intron variants. Three SNPs MA_10436040g0010_171180, MA_105586g0010_65505 and MA_10426383g0010_135796 were significantly associated *across* and *within* several traits, with all the modes of gene action being additive for the marker-trait interactions for the three SNPs (Table 2; Fig. 2).

**Table 2.**
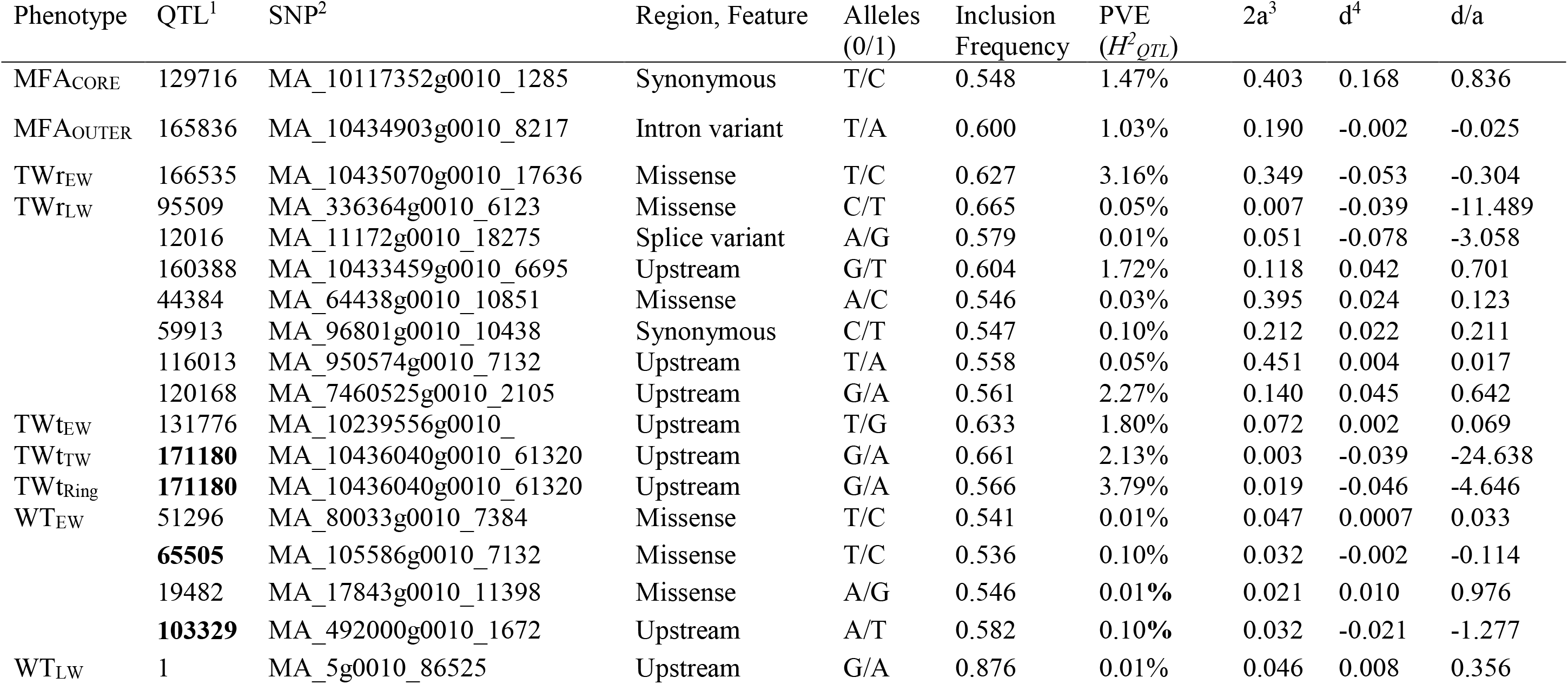
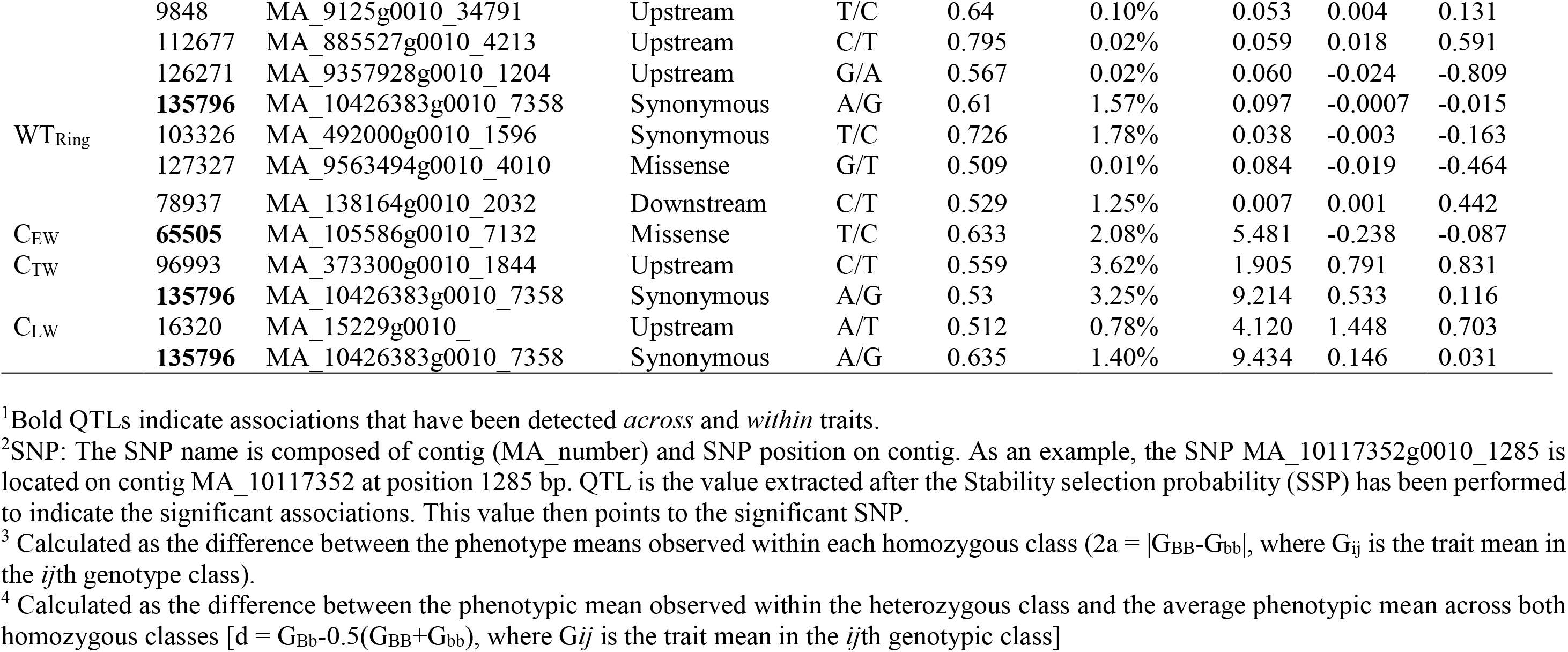
Phenotype, QTL position, allele frequencies and modes of allele inheritance.

**Fig. 2.**
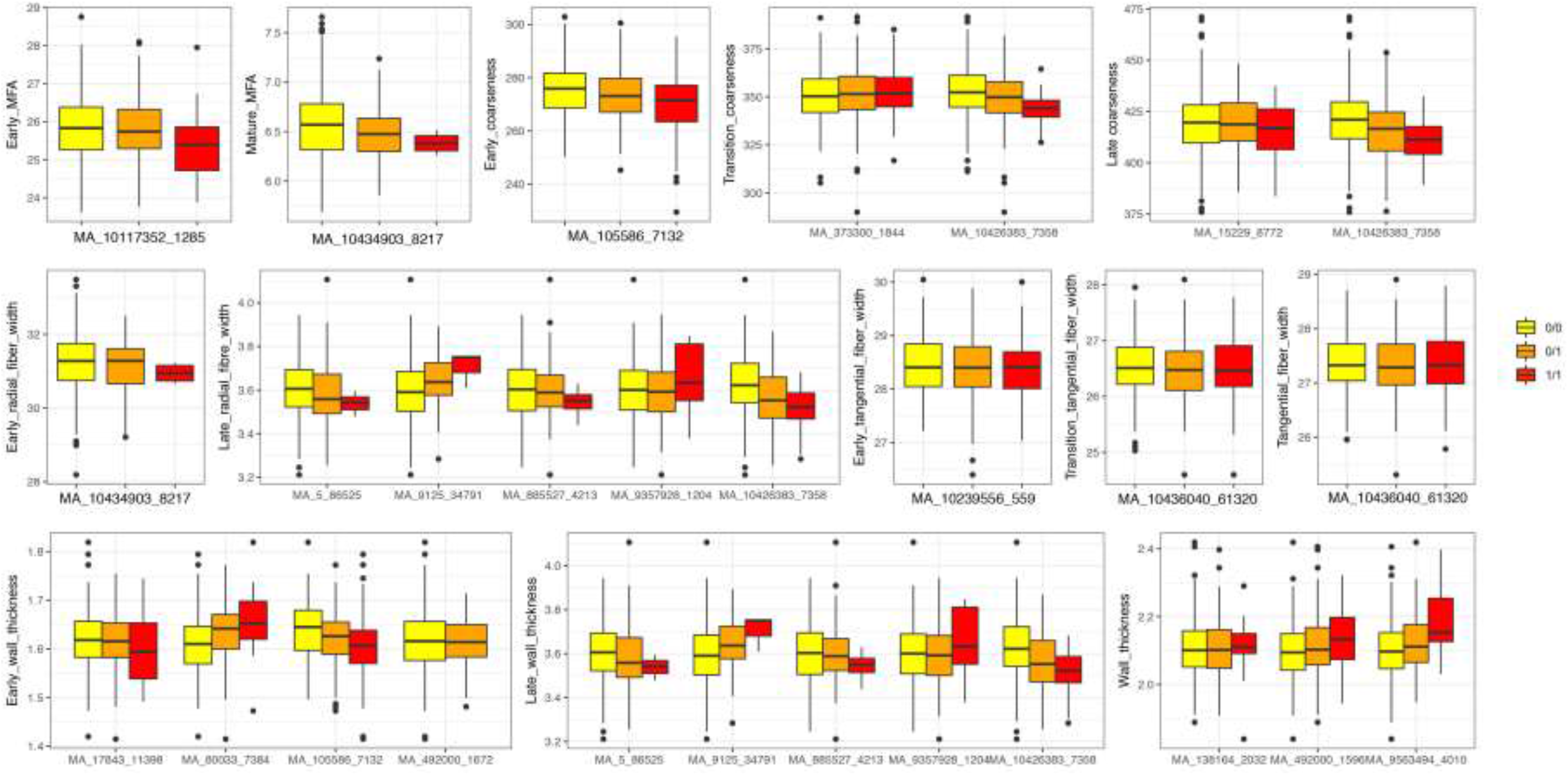
Box plot of the estimated genotypic effects for all significant associations identified in the study. The middle line represents the median value of the phenotype with that of the genotype. Upper and lower bounds of the box are the 25% (Q1) and 75% (Q3) quantile. Whiskers are Q1-1.5^*^Interquartile range (IQR) and Q3+1.5^*^IQR, therefore the outliers are values outside the range (Q1-1.5^*^IQR to Q3+1.5^*^IQ. Yellow, orange and red colored boxplots indicate the genotypic classes per SNP. Alleles (0/1) for all the SNPs have been defined in Table 2.

#### Genetic associations and genes of interest

Two associations for MFA were the intron variant MA_10434903g0010_8217 associated with MFA_OUTER_ explaining 1.03% of the PVE. MA_10117352g0010_1285 a synonymous variant explaining 1.47% PVE associated with MFA_OUTER_ occurred within gene MA_10117352g0010 homologous to Arabidopsis ONE HELIX PROTEIN (OHP). The gene is highly expressed especially in needles and shoots in spruce (Fig. 3). OHPs have been reported to be contitutively expressed and essential for photosynthesis in Arabidopsis, with mutants exhibiting severe growth defects (Beck *et al*., 2017).

Associations for radial tracheid widths were detected in earlywood and latewood. TWrEW was associated with a single missense SNP (MA_10435070g0010_17636) explaining 3.16% of the PVE and occurred within gene encoding nuclear transcription factor Y subunit A-7 (NF-YA7) (Table S1). NF-Y is a multimer complex binding CCAAT box in the promoter regions of many genes, and has multiple biological functions including growth regulation, cell size regulation, and responses to abiotic stresses (Zanetti *et al*., 2017; Zhao *et al*., 2017), including nitrogen deficiency in *Arabidopsis* (Sorin *et al*., 2014). The overexpression of the NF-YAs has been shown to stimulate growth during low nitrogen and phosphorous availability (Qu *et al*., 2015). This gene is highly and ubiquitously expressed in shoots and buds of spruce, indicating its important function in this species (Fig. 3).

**Fig. 3.**
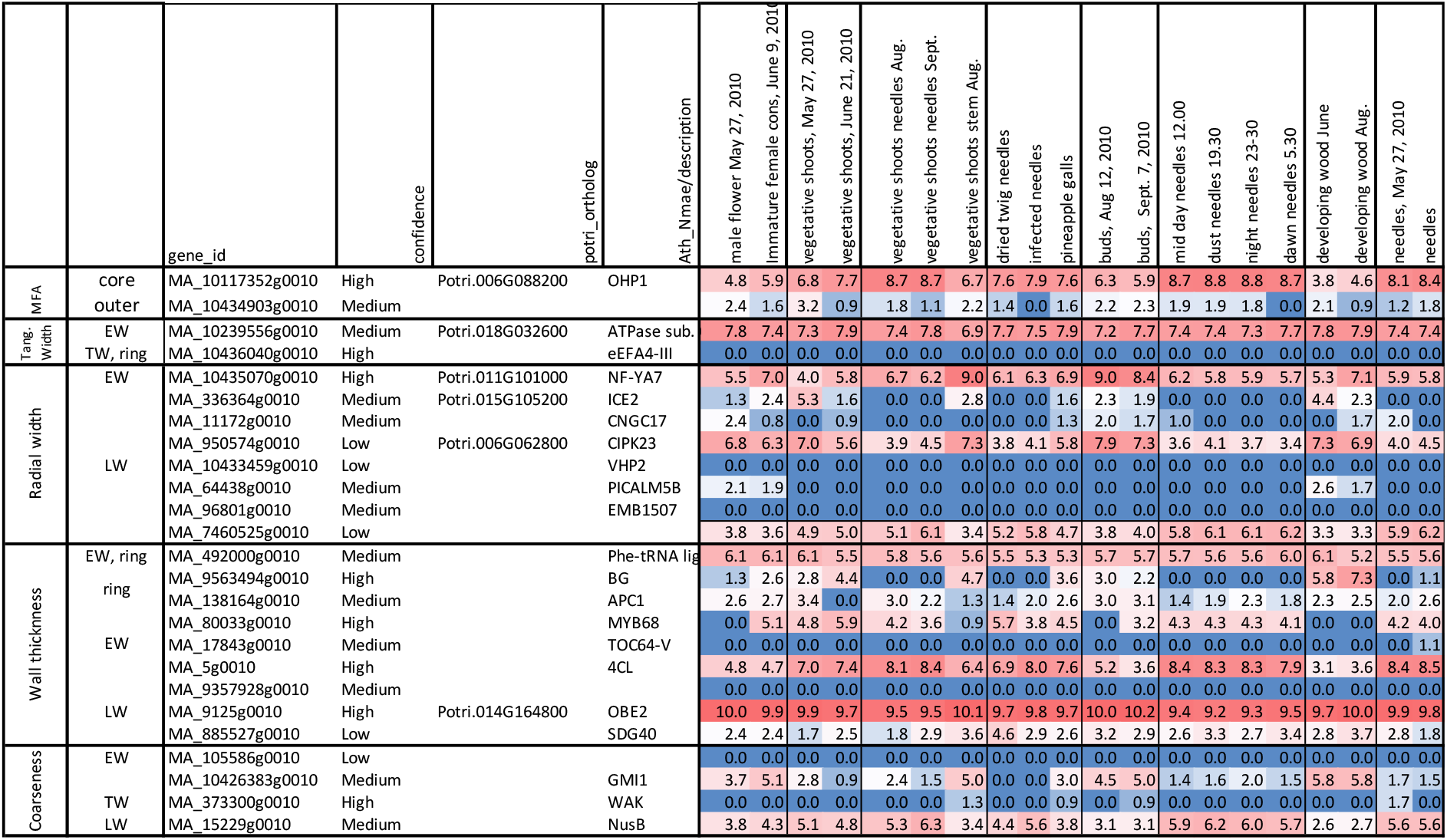
The heatmap showing the expression levels (VST values) of spruce candidate genes in different organs and tissues based on data of Nystedt et al., (2013) available at http://congenie.org.

TW_rLW_ with seven significant associations, had the highest number of significant associations per trait. Four of the associations were from modifier SNPs and the remaining three SNPs are ranked as moderate to low impact variants (Table 2). Two low impact missense SNPs MA_336364g0010_6123 and MA_64438g0010_10851 associated with TWrLW explained a small proportion of the PVE observed 0.01% and 0.03%, respectively. MA_336364g0010 is homologous to the *Arabidopsis INDUCER OF CBF EXPRESSION 2 (ICE2)* regulating deep-freezing tolerance by inducing *CBF1, CBF2* and *CBF3* genes (Table S1) (Kim *et al*., 2015). *CBF* genes have been identified to constitute a central node of hormone cross-talk during cold stress response and their expression is modulated by abscisic acid, gibberelins, jasmonate, ethylene and brassinosteroids (Barrero-Gil & Salinas, 2017). It has emerged that different hormone signaling pathways converge at the *CBF* promoter level, with the result of this hormone cross-talk being the fine-tuned transcript levels impacting on plant development and growth (Achard *et al*., 2008). In spruce, the gene is highly expressed in developing stems (Fig. 3) and strongly upregulated in the cambium and radial expansion zone (Fig. 4) supporting its role *in situ* in promoting the tracheid expansion. Since *CBFs* have already been identified as convergence points for hormones required for the regulation of plant growth under cold stress, these factors would warrant a detailed look in relation to their influence on wood tracheid development, especially during the time when the water stress and cold stress can be common. Gene MA_64438g0010 homologous to an *Arabidopsis PHOSPHATIDYLINOSITOL BINDING CLATHRIN ASSEMBLY PROTEIN 5B (PICALM5B),* a member of ENTH/ANTH/VHS superfamily (Table S1). The ENTH/ANTH/VHS superfamily is involved in clatrin assembly at secretory vescicles and is essential for vescicle intracellular trafficking and thus, cell growth and development (De Craene *et al*., 2012). The gene was observed expressed in developing wood (Fig. 4), indicating its importance for tracheid development in spruce. Indeed, genes of ENTH/ANTH/VHS family have been previously associated with secondary cell wall formation in *Populus* (Porth *et al*., 2013), and vescicle trafficking-related genes were seen upregulated coinciding with radial expansion of developing wood cells in aspen (Sundell *et al*., 2017). Such genes are therefore expected to be associated with tracheid radial expansion in spruce. Another gene associated with TWrLW was MA_950574g0010_7132, explaining a high PVE of 2.27%. It is remotely similar to *Arabidopsis CALCINEURIN-B-LIKE-INTERACTING SERINE/THREONINE-PROTEIN KINASE 23 (CBLPK23)* involved in regulation of HAK5-mediated high-affinity K^+^ uptake in calcium-dependent manner in *Arabidopsis* roots (Ragel *et al*., 2015). The confidence of the spruce model was low, but the gene was found highly expressed in developing shoots, buds and cones (Fig. 3), and during primary and secondary wall formation in developing spruce tracheids (Fig. 4) confirming that it was not a pseudogene. A *CALCINEURIN-B-LIKE* gene was found to explain the largest phenotypic variance in cell wall mannose content in white spruce (Beaulieu *et al*., 2011). These observations make the identified spruce *CBLPK23* gene an interesting candidate for calcium-dependent regulation of K^+^ uptake in developing tracheids, thus likely regulating tracheid expansion, similar to vessel element expansion, known to be dependent on K^+^ transport (Langer *et al*., 2002). Interestingly, there was another gene candidate related to K^+^ transport associated with tracheid radial width: the splice variant MA_11172g0010_18275 explaining 0.01% PVE (Table 2). This gene is homologous to *Arabidopsis CYCLIC NUCLEOTIDE-GATED CHANNEL 17 (CNGC17)* (Table S1). CNGCs are potassium channels involved in several plant physiological processes including root development, pollen tube growth and plant disease resistance (Ma *et al*., 2009). They regulate ion homeostasis within plants through the uptake of cations, which is essential for plant growth and development (Kaplan *et al*., 2007). Arabidopsis CNGC17 is localized in plasmamembrane and promotes protoplast expansion by regulating cation uptake (Ladwig *et al*., 2015). Its spruce homolog exhibited specific expression during latewood formation in August (Fig. 3), supporting its a role in latewood tracheid development.

Three significant associations were identified for tangential tracheid width components with an upstream variant MA_10436040g0010_61320 being detected across traits TWtTW and TWtRing (Table 2). This variant was detected on contig MA_10436040 with high inclusion frequencies explaining relatively high percentages of the variance observed, 2.13% for TWt_TW_ and 3.79% for TWt_Rin_ (Table 2). The associated gene - MA_10436040g0010 - is homologous to stress-related eukaryotic initiation factor 4A-III (eIF4A-IΠ) which also has a DEAD-box ATP-dependent RNA helicase 2, and is involved in RNA processing and nonsense-mediated mRNA decay in Arabidopsis, especially under hypoxia and heat stress (Table S1) (Pascuan *et al*., 2016). The spruce gene was not found expressed in available datasets (Figure 3; (Jokipii-Lukkari *et al*., 2017). SNP MA_10239556g0010_131776 was associated with TWtEW and explained a moderate amount of the PVE 1.80% (Table 2). The *Arabidopsis* homolog encodes a subunit C of a vacuolar ATP synthase, which is a membrane-bound enzyme complex/ion transporter that combines ATP synthesis and/or hydrolysis with the transport of protons across the tonoplast membrane. This gene was highly and ubiquitously expressed (Fig. 3; NorWood: http://norwood.congenie.org/norwood-v1.0/; Jokipii-Lukkari *et al*., 2017). All three SNPs were consistent with an additive mode of gene action (Table 2).

Twelve associations were detected for wall thickness components, with low to moderate *H^2^_QTL_* ranging from 0.01 to 1.78% (Table 2). Two of these associations (SNP MA_105586g0010_7132 and MA_10426383g0010_7358) were shared *across* cell wall thickness and coarseness traits.

Ring average for cell Wall Thickness (WT_Ring_) had three significant associations. The synonymous SNP MA_492000g0010_1672 had a high inclusion frequency (0.726) and explained the highest percentage of PVE (1.78%). The same SNP was associated with WT_EW_. The gene MA_492000g0010 is homologous to a tRNA synthetase beta subunit family protein, phenylalanyl-tRNA synthetase beta chain (Table S1). Consistent with its predicted general metabolic function in protein biosynthesis, it is highly and ubiquitously expressed in spruce tissues (Fig. 3). Missense SNP MA_9563494g0010_4010 and downstream variant MA_138164g0010_2032 explained 0.01% and 1.25% PVE, respectively. MA_9563494g0010_4010 is located on a gene *MA_9563494g0010* named as *Picea abies BIG GRAIN 2* (*PabBG2)* (Mishra *et al*., 2017) homologous to the *BIG GRAIN 1* gene (*OsaBG1*) in rice (Liu *et al*., 2015). *OsaBG1* encodes a membrane protein regulating auxin transport and sensitivity, and positively affecting plant biomass and seed size. The gene belongs to a small family containing nine members in spruce (Mishra et al., 2017). Auxin has long been known to act as a key hormone essential for the induction of vascular strands, cambial growth and secondary wall deposition (Uggla *et al*., 1998; Sundberg, 2000; Ranocha *et al*., 2013a; Ranocha *et al*., 2013b; Yang & Wang, 2016). *PabBG2* is highly expressed and specifically upregulated in the developing xylem (Fig. 3; Nystedt *et al*., 2013) with a peak of expression in the cambial zone (Fig. 6), coinciding with a peak of IAA distribution in wood forming tissues (Sundberg, 2000; Tuominen *et al*., 2000; Hellgren, 2003). It is therefore likely that *PabBG2* gene pays a major role in xylogenesis, as suggested by its association with tracheid cell wall thickness, and that it should be considered as main target for woody biomass increase. Moreover, the SNP MA_138164g0010_78937 explaining PVE 1.25% associated with WTRing was located on a gene homologous to the subunit of E3 ubiquitin complex encoded by *AtAPC1* and involved in cell cycle regulation by degradation of cyclin B1 (Guo *et al*., 2018). The E3 ubiquitin complex is also known in *Arabidopsis* to regulate auxin homeostasis (Gray *et al*., 1999; Kepinski & Leyser, 2004; Azpeitia & Alvarez-Buylla, 2012). Hence, the detection of two significant associations for WTRing that are potentially related to auxin regulation implies a close relation between auxin and cell wall thickness in spruce. A QTL in rice grain for width and weight, which are related to plant biomass, has been associated with a RING-type E3 ubiquitin ligase (Song *et al*., 2007). Several auxin responsive genes were also associated with tracheid width and MFA, which both are linked to cell wall thickness, in white spruce (Beaulieu *et al*., 2011).

WT_EW_ has three significant associations beside MA_492000g0010_103329 discussed above (Table 2). The missense variant MA_80033g0010_51296 was within a gene encoding a MYB transcription factor similar to Arabidopsis MYB68 (Table S1). This gene exhibited very low expression levels in the developing xylem but rather was expressed in young shoots and needles (Fig. 3). Different MYB transcription factors regulate plant developmental processes, and several have been identified as crucial factors for secondary wall deposition and lignification. Loblolly pine (*Pinus teada* L.) *PtMYB8* expressed in spruce induced secondary cell wall thickening (Bomal *et al*., 2008). White spruce (*P. glauca* L.) *PgMYB4* was associated with cell wall thickness and tracheid coarseness (Beaulieu *et al*., 2011), and has been shown to highly expressed during secondary cell wall formation and lignification in both white spruce and loblolly pine (Bedon *et al*., 2007). MYB encoded by *MA_80033g0010* could play a more indirect role in secondary wall regulation in spruce considering its expression (Fig. 3). Two remaining SNPs MA_17843g0010_19482 and MA_105586g0010_7132, had PVEs of 0.01% and 0.10%, respectively (Table 2). The former was a missense variant within a gene homologous to *Arabidopsis TOC64-V*. The latter was not matching any known gene and was also associated with C_EW_ and explaining a moderate percentage of PVE 2.08%. However, the two models were not expressed in any of the investigated organs (Fig. 3) suggesting that they may be the pseudogenes.

WT_LW_ was associated with four upstream variants and a single synonymous SNP MA_10426383g0010_7358. The four upstream variants explained PVE ranging from 0.01 to 0.10% whereas the synonymous SNP MA_10426383g0010_7358 having a high inclusion frequency explained a moderate amount of the PVE 1.57% (Table 2). *MA_10426383g0010* is homologous to *VIT_16s0098g01810* from *Vitis vinifera* (Table S1) annotated as encoding ATP binding protein that may be involved in chromosome organization and biogenesis (Davies & Coleman, 2000). *Arabidopsis* homolog - *GAMMA-IRRADIATION AND MITOMYCIN C INDUCED 1* (*GMI1*) is responsible for double strand repair via somatic homologous recombination (Böhmdorfer *et al*., 2011). The spruce gene shows increased expression in organs with active meristems (Fig. 3), which is expected for the function in DNA repair. The same SNP MA_10426383g0010_7358 was also associated to traits related to coarseness (C_TW_ and C_LW_) and explained a relatively high PVE 3.25% and 1.40%, respectively. It also had high inclusion frequencies for all three traits (WT_LW_, C_TW_ and C_LW_) (Table 2). The associated gene might therefore be a good candidate to explore for effects on tracheid development, especially since it is highly expressed in the developing wood (Fig. 3; Nystedt *et al*., 2013). SNP MA_5g0010_1 associated with WT_LW_ was detected upstream of gene MA_5g0010 homologous to the gene of *Populus trichocarpa* annotated as encoding 4-coumarate-CoA ligase (4CL) a key enzyme in monolignol biosynthetic pathway. However, the Arabidopsis homolog of *MA_5g0010* is annotated as fatty acyl CoA synthase involved in lipid biosynthesis. *MA_5g0010* is not expressed in the developing wood but it is highly expressed in young vegetative shoots and needles, including the infected needles (Fig. 3), making it unlikely candidate for lignin biosynthesis gene in developing wood but suggesting a rather indirect function in regulation of tracheid cell wall thickness. The SNP MA_9125g0010_34791 associated with WT_LW_ was located upstream of a gene homologous to *Arabidopsis OBERON2 (OBE2)* encoding a plant homeodomain (PHD) finger protein (Table S1) (Lee *et al*., 2009). Homeodomain genes encode transcription factors central in the regulation of plant developmental processes (Chew *et al*., 2013). *OBE1* and *OBE2* redundanlty regulate meristem establishment and maintenance in *Arabidopsis* (Saiga *et al*., 2008). The spruce *OBE2* gene is highly and ubiquitously expressed in vegetative and reproductive organs (Fig. 3) including developing wood where it shows high expression during secondary wall deposition (Fig. 4) and therefore it could have a direct role in cell wall thickening in tracheids. SNP MA_885527g0010_112677 associated with WT_LW_ was found upstream of a gene containing a SET domain. SET domain proteins have been identified in *Arabidopsis* to be involved in the epigenetic control of genes involved in a wide range of activities including plant growth (Thorstensen *et al*., 2011). A link has also been established between PHD finger proteins and SET domain proteins in the regulation of developmental transitions in *Arabidopsis* where PHD finger proteins VEL1, VRN5 and VIN3 interacting with H3K27me3 repress FLC transcription allowing for the transition from vegetative to reproductive development (Thorstensen *et al*., 2011). *MA_885527g0010* is highly upregulated in developing wood sample from August that is involved in latewood biosynthesis (Fig. 3) suggesting its direct role in latewood tracheid development.

**Fig 4.**
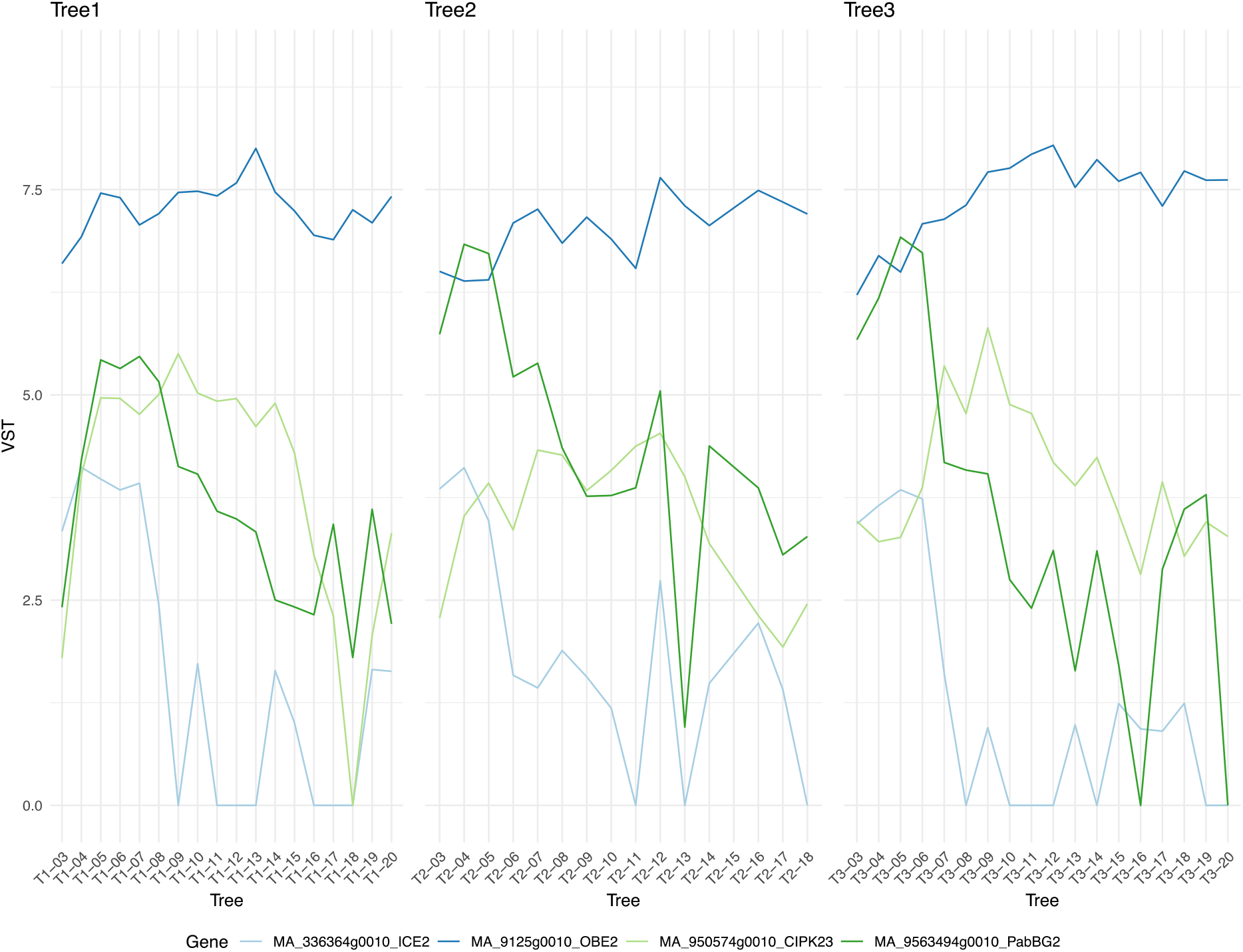
Expression profiles of selected candidate genes in wood developing tissues of spruce based on Norwood dataset (NorWood: http://norwood.congenie.org/norwood-v1.0/; Jokipii-Lukkari *et al*., 2017). *MA_336364g0010 (ICE2)* gene and *MA_9563494g0010* – *PabBG2* have high expression in the cambium coinciding with a peak of IAA distribution, whereas *MA_950574g0010 (CIPK23)* and *MA_9125g0010 (OBE2*) have broad expression pattern with high levels observed during primary and secondary wall formation.

A total of five significant associations were identified for coarseness traits explaining moderate to high *H^2^_QTL_* ranging from 0.78 to 3.62% (Table 2). Three of them concerning SNP MA_105586g0010_7132 and MA_10426383g0010_7358 were also associated with WT_EW_ and WT_LW_, and discussed above. Upstream variant MA_373300g0010_1844 associated with C_TW_ explained a relatively high percentage of PVE 3.62% and was consistent with a partial to fully dominant mode of gene action (Table 2) as shown by the genotypic effects (Fig 2). The gene is similar to POPTR_0005s15330g from *Populus trichocarpa* annotated as a wall-associated receptor kinase (WAK). WAKs have been previously reported to be associated with average ring width and the proportion of earlywood in white spruce (Beaulieu *et al*., 2011) and with MFA in *Populus* (Porth *et al*., 2013). The gene is expressed primarily in early season needles and late season stem from vegetative shoots but there is no detectable expression in developing xylem (Fig. 3; Nystedt et al., 2013), suggesting its indirect involvement regulation of in tracheid coarseness.

This work presents the first genome wide dissection of wood tracheid traits in Norway spruce. A total of 30 significant associations were detected for all investigated traits. These associations have identified a set of genes that could be exploited to alter wood tracheid traits for improving solid wood properties for its use in industrial processes. Previous studies utilizing a LASSO penalized analysis approach were limited in the nature and number of molecular markers available (Ma *et al*., 2002; Li *et al*., 2014), with our study representing a major advance by using 178101 SNPs with a functional mapping approach. The relatively small number of associations is comparable to other association studies of complex growth traits in forest trees, were a few associations are detected with a relatively small proportion of the genetic variation being explained (Cappa *et al*., 2013; Porth *et al*., 2013; McKown *et al*., 2014; Allwright *et al*., 2016; Lamara *et al*., 2016). It can be argued that many of the alleles causing variation for polygenic traits may be either rare or have small effects and current GWAS methods lack the power to detect them, thus the small number of significant associations (Thornton *et al*., 2013; De La Torre *et al*., 2019). The small number of associations being reported could also be largely due to the small sample sizes in these studies for such complex traits. The statistical power required to detect associations between molecular markers and a trait is heavily dependent upon the sample size (Visscher *et al*., 2017; Müller *et al*., 2018). Due to the challenges of developing large populations for GWAS in conifers, the majority of the studies utilize a few hundred individuals from natural populations, which limit the statistical power for GWAS. Potential viable options are being developed which utilize a combination of information from multiple populations using Meta-GWAS and Joint-GWAS (Mägi & Morris, 2010; Bernal Rubio *et al*., 2015). These approaches are now being applied in some recent forest tree studies (Müller *et al*., 2018) and could be the next level of analysis using our application of latent traits on these complex traits.

#### Data Availability

All the latent traits, genotypic data, SNP position files the association mapping scripts used for the analysis are publicly available at are available from zenodo.org at https://doi.org/10.5281/zenodo.1480536 (DOI: 10.5281/zenodo.1480536). Raw sequence data for all the samples utilized in the study are found through the European Nucleotide Archive under accession number PRJEB29652. The Norway spruce genome assemblies and resources are available from http://congenie.org/pabiesgenome.

## Supporting information

Candidate genes

## Acknowledgements

We acknowledge the Bio4Energy consortium for giving us accesss to the Silviscan wood properties data collection. We also acknowledge the support from Science for Life Laboratory, the Knut and Alice Wallenberg Foundation, the National Genomics Infrastructure funded by the Swedish Research Council, and Uppsala Multidisciplinary Center for Advanced Computational Science for assistance with massively parallel sequencing and access to the UPPMAX computational infrastructure. John Baison was supported though a postdoc position funded by the Kempe foundation.

