## Supplementary material for "Genetic control of tracheid properties in Norway spruce wood": Candidate genes

### *New Phytologist* Supporting Information

Article title: **Genetic control of tracheid properties in Norway spruce wood**

Article acceptance date: Click here to enter a date.

The following Supporting Information is available for this article:

**Table S1.** BLAST search results from contigs with significant QTL from spline model

|  | Trait | | Marker | *Contig* | | *Putative Genes* |
| --- | --- | --- | --- | --- | --- | --- |
|  | MFA_OUTER_ | | 129716 | MA_10117352g0010_129716 | | - Arabidopsis ONE HELIX PROTEIN 1 (OHP1) Chlorophyll a-b binding protein. Are expressed constitutively in green plant tissues and their levels increase in response to light stress.  Ath: [AT5G02120.1](http://atgenie.org/transcript?id=AT5G02120.1)  Potri: [Potri.006G088200.1](http://popgenie.org/transcript?id=Potri.006G088200.1)  [GO:0003674](http://amigo.geneontology.org/cgi-bin/amigo/term_details?term=GO:0003674): function  [GO:0009507](http://amigo.geneontology.org/cgi-bin/amigo/term_details?term=GO:0009507): chloroplast  [GO:0009535](http://amigo.geneontology.org/cgi-bin/amigo/term_details?term=GO:0009535): thylakoid membrane (sensu Viridiplantae)  [GO:0009644](http://amigo.geneontology.org/cgi-bin/amigo/term_details?term=GO:0009644): response to high light intensity |
|  | MFA_CORE_ | | 165836 | MA_10434903g0010_165836 | | Uncharacterized |
|  | FWr_EW_ | | 166535 | MA_10435070g0010_166535 | | -NF-YA7 protein of Arabidopsis. CCAAT-binding factor which is a heteromeric transcription factor that consists of two different components both needed for DNA-binding  Ath: AT1G30500.1  Potri: [Potri.011G101000.1](http://popgenie.org/transcript?id=Potri.011G101000.1)  [GO:0003700](http://amigo.geneontology.org/cgi-bin/amigo/term_details?term=GO:0003700): transcription factor activity  [GO:0005634](http://amigo.geneontology.org/cgi-bin/amigo/term_details?term=GO:0005634): nucleus  [GO:0006355](http://amigo.geneontology.org/cgi-bin/amigo/term_details?term=GO:0006355): regulation of transcription, DNA-dependent |
|  | FWr_LW_ | | 95509 | MA_336364g0010_95509 | | -Inducer of CBF expression 2 (ICE2). This protein has the basic helix-loop-helix (bHLH) is a [protein](http://en.wikipedia.org/wiki/Protein) [structural motif](http://en.wikipedia.org/wiki/Structural_motif) that characterizes a family of [transcription factors](http://en.wikipedia.org/wiki/Transcription_factors). bHLH transcription factors are often important in development or cell activity. ICE2 is involved in freezing adaptation in Arabidopsis.  Ath: AT1G12860.1  Potri: [Potri.015G105200.1](http://popgenie.org/transcript?id=Potri.015G105200.1)  [GO:0006355](http://amigo.geneontology.org/cgi-bin/amigo/term_details?term=GO:0006355): regulation of transcription, DNA-dependent  [GO:0030528](http://amigo.geneontology.org/cgi-bin/amigo/term_details?term=GO:0030528): transcription regulator activity |
|  |  |  | 12016 | MA_11172g0010_12016 | | -Cyclic nucleotide-gated channel 17 (CNGC17) of Arabidopsis. There are commonly additional regulatory domains, which serve to regulate ion conduction and channel gating. The pores may also be homotetramers or hetrotetramers; where hetrotetramers may be encoded as distinct genes or as multiple pore domains within a single polypepetide.  Ath: AT4G30360.1  Potri: Potri.018G009200.1  [GO:0005216](http://amigo.geneontology.org/cgi-bin/amigo/term_details?term=GO:0005216): ion channel activity  [GO:0016020](http://amigo.geneontology.org/cgi-bin/amigo/term_details?term=GO:0016020): membrane  [GO:0055085](http://amigo.geneontology.org/cgi-bin/amigo/term_details?term=GO:0055085): transmembrane transport |
|  |  |  | 160388 | MA_10433459g0010_160388 | | -BnaCnng64390D protein from *Brassica napus* characterized as an inorganic diphosphatase. This enzyme belongs to the family of [hydrolases](http://en.wikipedia.org/wiki/Hydrolase) acting on acid anhydrides in phosphorus-containing anhydrides. Arabidopsis vacuolar H+ pyrophosphatase 2 (VHP2)  [GO:0004427](https://www.ebi.ac.uk/QuickGO/term/GO:0004427): inorganic diphosphatase activity  [GO:0009678](https://www.ebi.ac.uk/QuickGO/term/GO:0009678): hydrogen-translocating pyrophosphatase activity |
|  |  | | 44384 | MA_64438g0010_44384 | | -ENTH/ANTH/VHS superfamily protein. Membrane trafficking involves the complex regulation of proteins and lipids intracellular localization and is required for metabolic uptake, cell growth and development. Arabidopsis PHOSPHATIDYLINOSITOL BINDING CLATHRIN ASSEMBLY PROTEIN 5B, (PICALM5B)  Ath: AT4G02650.1  Potri: [Potri.019G063700.1](http://popgenie.org/transcript?id=Potri.019G063700.1)  [GO:0005488](http://amigo.geneontology.org/cgi-bin/amigo/term_details?term=GO:0005488): binding  [GO:0005543](http://amigo.geneontology.org/cgi-bin/amigo/term_details?term=GO:0005543): phospholipid binding  [GO:0030118](http://amigo.geneontology.org/cgi-bin/amigo/term_details?term=GO:0030118): clathrin coat  [GO:0048268](http://amigo.geneontology.org/cgi-bin/amigo/term_details?term=GO:0048268): clathrin cage assembly |
|  |  | | 59913 | MA_96801g0010_59913 | | - U5 small nuclear ribonucleoprotein helicase. Plays an essential role in pre-mRNA splicing as component of the U5 snRNP and U4/U6-U5 tri-snRNP complexes. Involved in spliceosome assembly, activation and disassembly. Mediates changes in the dynamic network of RNA-RNA interactions in the spliceosome. Arabidopsis *embryo defective 1507* (*EMB 1507*)  Ath: AT1G20960.2  Potri: [Potri.015G095500.1](http://popgenie.org/transcript?id=Potri.015G095500.1)  [GO:0003676](http://amigo.geneontology.org/cgi-bin/amigo/term_details?term=GO:0003676): nucleic acid binding  [GO:0003677](http://amigo.geneontology.org/cgi-bin/amigo/term_details?term=GO:0003677) DNA binding  [GO:0004386](http://amigo.geneontology.org/cgi-bin/amigo/term_details?term=GO:0004386): helicase activity  [GO:0016787](http://amigo.geneontology.org/cgi-bin/amigo/term_details?term=GO:0016787): hydrolase activity |
|  |  | | 116013 | MA_950574g0010_116013 | | - CBL-interacting serine/threonine-protein kinase 23 (CIPK23). Regulates HAK5-Mediated High-Affinity K+ Uptake in Arabidopsis Roots  Beta vulgaris: [XM_010688807.2](https://www.ncbi.nlm.nih.gov/nucleotide/XM_010688807?report=genbank&log$=nuclalign&blast_rank=3&RID=36YNER2K014) |
|  | FWt_EW_ | | 131776 | MA_10239556g0010_131776 | | -ATP synthase subunit C is membrane-bound enzyme complexes/ion transporters that combine ATP synthesis and/or hydrolysis with the transport of protons across a membrane.  Ath: AT2G25610.1  Potri: [Potri.018G032600.1](http://popgenie.org/transcript?id=Potri.018G032600.1)  [GO:0015078](http://amigo.geneontology.org/cgi-bin/amigo/term_details?term=GO:0015078): hydrogen ion transporter activity  [GO:0015991](http://amigo.geneontology.org/cgi-bin/amigo/term_details?term=GO:0015991): ATP hydrolysis coupled proton transport  [GO:0033177](http://amigo.geneontology.org/cgi-bin/amigo/term_details?term=GO:0033177): proton-transporting two-sector ATPase complex, proton-transporting domain |
|  | FWt_TW_ | | **171180** | MA_10436040g0010_**171180** | | - Eukaryotic initiation factor 4A-III (eIF4A-III) containing DEAD-box ATP-dependent RNA helicase 2. The DEAD-box RNA helicase family comprises enzymes that participate in RNA metabolism.  Ath: [AT3G19760.1](http://atgenie.org/transcript?id=AT3G19760.1)  Potri: [Potri.007G070000.1](http://popgenie.org/transcript?id=Potri.007G070000.1)  [GO:0003676](http://amigo.geneontology.org/cgi-bin/amigo/term_details?term=GO:0003676): nucleic acid binding  [GO:0004386](http://amigo.geneontology.org/cgi-bin/amigo/term_details?term=GO:0004386): helicase activity  [GO:0008026](http://amigo.geneontology.org/cgi-bin/amigo/term_details?term=GO:0008026): ATP dependent helicase activity |
|  | FWt_Ring_ | | **171180** | MA_10436040g0010_**171180** | | -DEAD-box ATP-dependent RNA helicase 2. The DEAD-box RNA helicase family comprises enzymes that participate in every aspect of RNA metabolism. Eukaryotic initiation factor 4A-3 also associated with the contig.  Ath: [AT3G19760.1](http://atgenie.org/transcript?id=AT3G19760.1)  Potri: [Potri.007G070000.1](http://popgenie.org/transcript?id=Potri.007G070000.1)  [GO:0003676](http://amigo.geneontology.org/cgi-bin/amigo/term_details?term=GO:0003676): nucleic acid binding  [GO:0004386](http://amigo.geneontology.org/cgi-bin/amigo/term_details?term=GO:0004386): helicase activity  [GO:0008026](http://amigo.geneontology.org/cgi-bin/amigo/term_details?term=GO:0008026): ATP dependent helicase activity |
| WT_EW_ | | 51296 | | | MA_80033g0010_51296 | -MYB family transcription factor. Gene JCGZ_12585 from Jatropha curcas (Barbados nut). Arabidopsis MYB68.  Ath: [AT5G65790.1](http://atgenie.org/transcript?id=AT5G65790.1)  Potri: [Potri.018G095900.1](http://popgenie.org/transcript?id=Potri.018G095900.1) |
|  | | 19482 | | | MA_17843g0010_19482 | **-** Arabidopsis translocon at the outer membrane of chloroplasts 64-V (TOC64-V)  **Ath:** [AT5G09420](http://atgenie.org/gene?id=AT5G09420)  **Potri:** [Potri.001G205300](http://popgenie.org/gene?id=Potri.001G205300)  [GO:0005515](http://amigo.geneontology.org/cgi-bin/amigo/term_details?term=GO:0005515): protein binding  [GO:0016884](http://amigo.geneontology.org/cgi-bin/amigo/term_details?term=GO:0016884): carbon-nitrogen ligase activity, with glutamine as amido-N-donor |
|  | | 65505 | | | MA_105586g0010 | **Uncharacterized** |
|  | | 103329 | | | MA_492000g0010_103329 | **-** tRNA synthetase beta subunit family protein. phenylalanyl-tRNA synthetase beta chain  **Ath:** [AT1G72550](http://atgenie.org/gene?id=AT1G72550).1  **Potri:** [Potri.013G068100.1](http://popgenie.org/transcript?id=Potri.013G068100.1)  [GO:0000287](http://amigo.geneontology.org/cgi-bin/amigo/term_details?term=GO:0000287): magnesium ion binding  [GO:0003723](http://amigo.geneontology.org/cgi-bin/amigo/term_details?term=GO:0003723): RNA binding  [GO:0004826](http://amigo.geneontology.org/cgi-bin/amigo/term_details?term=GO:0004826): phenylalanine-tRNA ligase activity  [GO:0006412](http://amigo.geneontology.org/cgi-bin/amigo/term_details?term=GO:0006412): protein biosynthesis  [GO:0006432](http://amigo.geneontology.org/cgi-bin/amigo/term_details?term=GO:0006432): phenylalanyl-tRNA aminoacylation |
|  | WT_LW_ | | 1 | MA_5g0010_1 | | -Putative 4-coumarate-CoA ligase from Picea glauca (White spruce), Arabidopsis fatty acyl CoA synthase  - in the coniferous gymnosperm Pinus radiata substantially affected plant phenotype and resulted in dwarfed plants with a “bonsai tree-like” appearance.  Ath: [AT4G05160.1](http://atgenie.org/transcript?id=AT4G05160.1)  Potri: [Potri.017G112800.1](http://popgenie.org/transcript?id=Potri.017G112800.1)  [GO:0003824](http://amigo.geneontology.org/cgi-bin/amigo/term_details?term=GO:0003824): catalytic activity  [GO:0008152](http://amigo.geneontology.org/cgi-bin/amigo/term_details?term=GO:0008152): metabolism  [GO:0004321](http://amigo.geneontology.org/cgi-bin/amigo/term_details?term=GO:0004321): fatty-acyl-CoA synthase activity  [GO:0005777](http://amigo.geneontology.org/cgi-bin/amigo/term_details?term=GO:0005777): peroxisome  [GO:0009695](http://amigo.geneontology.org/cgi-bin/amigo/term_details?term=GO:0009695): jasmonic acid biosynthesis  [GO:0016207](http://amigo.geneontology.org/cgi-bin/amigo/term_details?term=GO:0016207): 4-coumarate-CoA ligase activity |
|  |  |  | 9848 | MA_9125g0010_9848 | | -plant homeodomain finger protein similar to, Arabidopsis Oberon2 (OBE2) responsible for maintenance and establishment of root and shoot meristems. maintenance and/or establishment of both the shoot and root apical meristems in Arabidopsis.  Ath: [AT5G48160.2](http://atgenie.org/transcript?id=AT5G48160.2)  Potri: [Potri.014G164800.2](http://popgenie.org/transcript?id=Potri.014G164800.2)  [GO:0005515](http://amigo.geneontology.org/cgi-bin/amigo/term_details?term=GO:0005515): protein binding  [GO:0005634](http://amigo.geneontology.org/cgi-bin/amigo/term_details?term=GO:0005634): nucleus  [GO:0008270](http://amigo.geneontology.org/cgi-bin/amigo/term_details?term=GO:0008270): zinc ion binding  [GO:0009793](http://amigo.geneontology.org/cgi-bin/amigo/term_details?term=GO:0009793): embryonic development (sensu Magnoliophyta)  [GO:0010071](http://amigo.geneontology.org/cgi-bin/amigo/term_details?term=GO:0010071): root meristem specification  [GO:0010078](http://amigo.geneontology.org/cgi-bin/amigo/term_details?term=GO:0010078): maintenance of root meristem identity  [GO:0010468](http://amigo.geneontology.org/cgi-bin/amigo/term_details?term=GO:0010468): regulation of gene expression |
|  |  |  | 112677 | MA_885527g0010_112677 | | -SET domain (It has been demonstrated that association of SET domain and myotubularin-related [proteins](http://en.wikipedia.org/wiki/Protein) modulates [growth](http://en.wikipedia.org/wiki/Cell_growth) control) and Rubisco LSMT (They allow binding of the protein to substrate, such as the N-terminal tails of histones H3 and H4 and the large subunit of the Rubisco holoenzyme complex) domains. These two domains interact in lysine methylation. Arabidopsis homolog encodes SET domain group 40 protein (SDG40).  Ath:   [AT5G17240.1](http://atgenie.org/transcript?id=AT5G17240.1)  Potri: [Potri.004G081400.2](http://popgenie.org/transcript?id=Potri.004G081400.2) |
|  |  |  | 126271 | MA_9357928g0010_126271 | | -the contig has RRM motif is probably diagnostic of an RNA binding protein. RRMs are found in a variety of RNA binding proteins, including various hnRNP proteins, proteins implicated in regulation of alternative splicing, and protein components of snRNPs.  Ath: AT5G46870.1  Potri: [Potri.001G138400.1](http://popgenie.org/transcript?id=Potri.001G138400.1)  [GO:0003676](http://amigo.geneontology.org/cgi-bin/amigo/term_details?term=GO:0003676): nucleic acid binding  [GO:0016491](http://amigo.geneontology.org/cgi-bin/amigo/term_details?term=GO:0016491): oxidoreductase activity  [GO:0055114](http://amigo.geneontology.org/cgi-bin/amigo/term_details?term=GO:0055114): oxidation reduction |
|  |  |  | **135796** | MA_10426383g0010_**135796** | | -Gene VIT_16s0098g01810. An ATP binding protein. Involved in chromosome organization and biogenesis. From Vitis vinifera (Grape). Arabidopsis GAMMA-IRRADIATION AND MITOMYCIN C INDUCED 1 (GMI1) is involved in double strand break repair.  Ath:  [AT5G24280.1](http://atgenie.org/transcript?id=AT5G24280.1)  Potri: [Potri.012G016100.1](http://popgenie.org/transcript?id=Potri.012G016100.1) |
|  | WT_Ring_ | | 103326 | MA_492000g0010_103326 | | -Phenylalanyl-tRNA synthetase beta chain, putative. Ath: AT1G72550.1 Potri: [Potri.013G068100.1](http://popgenie.org/transcript?id=Potri.013G068100.1) [GO:0000287](http://amigo.geneontology.org/cgi-bin/amigo/term_details?term=GO:0000287): magnesium ion binding [GO:0003723](http://amigo.geneontology.org/cgi-bin/amigo/term_details?term=GO:0003723): RNA binding [GO:0004826](http://amigo.geneontology.org/cgi-bin/amigo/term_details?term=GO:0004826): phenylalanine-tRNA ligase activity [GO:0005524](http://amigo.geneontology.org/cgi-bin/amigo/term_details?term=GO:0005524): ATP binding [GO:0006412](http://amigo.geneontology.org/cgi-bin/amigo/term_details?term=GO:0006412): protein biosynthesis [GO:0006432](http://amigo.geneontology.org/cgi-bin/amigo/term_details?term=GO:0006432): phenylalanyl-tRNA aminoacylation |
|  |  |  | 127327 | MA_9563494g0010_127327 | | -Encodes *Picea abies* BIG GRAIN 2 (*Pab*BG2). BIG GRAIN (BG) proteins are found in all plants and spruce has 9 BG genes and one BG-like gene. Rice BG1 is a positive regulator of biomass and seed size acting via auxin signaling.  Ath: At1g54200 |
|  |  | | 78937 | MA_138164g0010_78937 | | - E3 ubiquitin ligase. Ubiquitination cascade is one of the main pathways of post-translational regulation of gene expression. Arabidopsis homolog encodes ANAPHASE PROMOTING COMPLEX 1 (APC1) functioning in cyclin B1 homeostasis in embryo- and gametogenesis.  Ath: [AT5G05560](http://atgenie.org/gene?id=AT5G05560).1  Potri: [Potri.008G070300.1](http://popgenie.org/transcript?id=Potri.008G070300.1)  [GO:0000151](http://amigo.geneontology.org/cgi-bin/amigo/term_details?term=GO:0000151): ubiquitin ligase complex  [GO:0004842](http://amigo.geneontology.org/cgi-bin/amigo/term_details?term=GO:0004842): ubiquitin-protein ligase activity  [GO:0006511](http://amigo.geneontology.org/cgi-bin/amigo/term_details?term=GO:0006511): ubiquitin-dependent protein catabolism |
|  | C_EW_ | | **65505** | MA_105586g0010_**65505** | | -Neurabin-1-like protein. Contains a PDZ (Proteins containing PDZ domains play a key role in anchoring receptor proteins in the membrane to cytoskeletal components.) and SAM domains (SAM as a protein interaction domain involved in developmental regulation). |
|  | C_TW_ | | 96993 | MA_373300g0010_96993 | | -Gene POPTR_0005s15330g. Wall-associated receptor kinase C-terminal. From Populus trichocarpa (Western balsam poplar)  Potri: [Potri.T064000.1](http://popgenie.org/transcript?id=Potri.T064000.1)  [GO:0004672](http://amigo.geneontology.org/cgi-bin/amigo/term_details?term=GO:0004672): protein kinase activity  [GO:0005524](http://amigo.geneontology.org/cgi-bin/amigo/term_details?term=GO:0005524): ATP binding  [GO:0006468](http://amigo.geneontology.org/cgi-bin/amigo/term_details?term=GO:0006468): protein amino acid phosphorylation |
|  | C_LW_ | | 16320 | MA_15229g0010_16320 | | -Putative uncharacterized protein from Picea sitchensis (Sitka spruce). Involved in regulation of transcription.  -pfam domain NusB protein is involved in the regulation of rRNA biosynthesis by transcriptional antitermination.  [GO:0003723](http://amigo.geneontology.org/cgi-bin/amigo/term_details?term=GO:0003723): RNA binding  [GO:0006355](http://amigo.geneontology.org/cgi-bin/amigo/term_details?term=GO:0006355): regulation of transcription, DNA-dependent |
|  |  |  | **135796** | MA_10426383g0010_**135796** | | -Arabidopsis GAMMA-IRRADIATION AND MITOMYCIN C INDUCED 1 (GMI1)  -Gene VIT_16s0098g01810. A ATP binding protein. Involved in chromosome organization and biogenesis. Vitis vinifera (Grape). |

**Methods S1** Quadratic spline

Quadratic spline with multiple knots:

*β(t) = b_0_ + b_1_t + b_2_(t-t_t_)^2^_+_ + b_3_(t-t_2_)_+_+…+ b_1+k_(t-t_k_),* (1)

where *t*_i_ (*i*=1,…,*k*; *t*_1_<*t*_2_…<*t_k_*) are defined as knots, and *(t – t_i_)_+_ = (t – t_i_)* if *t > t_i_* (*t_i_*>0; *i*=1,…,*k*), and otherwise is equal to zero. Generally speaking, the location and amount of knots have to be properly defined in order to provide an accurate description of the data under investigation (Li *et al.*, 2015). In our case, since the growth pattern of wood property traits were not complex, we choose two knots of the time interval.

Hence, the quadratic spline model to describe the growth trajectory of individual *i* applied in this study was defined as:

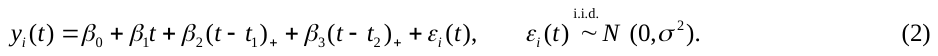

Then the intercept*β*_0_, slope*β*_1_,*β*_2_ (Knot 1 (k1)) and *β*_3_ (Knot 2 (k2)) are estimated by standard least squares, and their estimates were considered as the latent trait in the subsequent QTL analysis conducted in R-studio (Team, 2015).

**Methods S2** The LASSO model

The LASSO model:

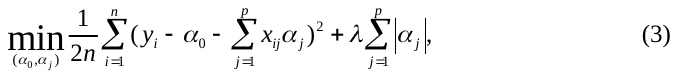

where *y_i_* is the phenotypic value of individual *i* (*i*=1,…,*n*; *n* is the total number of individuals) of one of the latent trait (*β*_0_, *β*_1_, *β*_2_, *β*_3_), *x_ij_* is the genotypic value of individual *i* and marker *j* coded as 0, 1 and 2 for three marker genotypes AA, AB and BB, respectively, *α*_0_ is the population mean parameter, *α_j_* is the effect of marker *j* (*i*=1,…,*n*; *n* is the total number of markers), and *λ* (>0) is a shrinkage tuning parameter. A fundamental idea of LASSO is to utilize the penality function
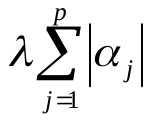
to shrink the SNP effects toward zero, and only keep a small number of important SNPs which are highly associated with the trait in the model.

**Methods S3** PVE evaluation of a QTL

PVE evaluation:

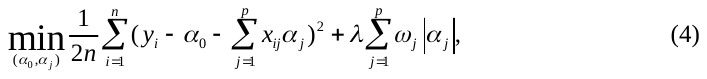

where the penalty function has a separate weight
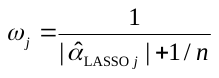
 (
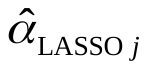
 represents the initial LASSO estimates from (3)) for each independent SNP, and the rest of the parameters are defined in the same way as in the standard LASSO regression (3). By the aid of the SNP specific tuning factor, adaptive LASSO should provide un-biased estimates to the effect size and PVE of a QTL.

**Methods S4** Genotypic Effect Calculation

The line in the middle is the median value of the phenotype with that genotype. upper bound and lower bound of the box is the 25% (Q1) and 75% (Q3) quantile. Whiskers are Q1-1.5*IQR and Q3+1.5*IQR. So the outliers are values outside the range (Q1-1.5*IQR,Q3+1.5*IQR)

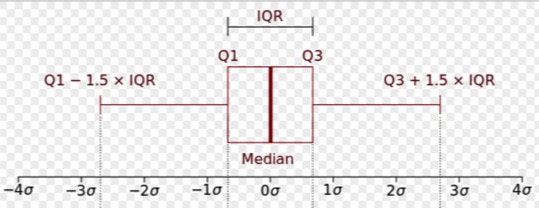
